# Dysbiosis in a canine model of human fistulizing Crohn’s disease

**DOI:** 10.1101/815589

**Authors:** Ana Maldonado-Contreras, Lluís Ferrer, Caitlin Cawley, Sarah Crain, Juan Toscano, Doyle V. Ward, Andrew Hoffman

**Affiliations:** Department of Microbiology and Physiological Systems, University of Massachusetts Medical School, Worcester, MA, 01655; Cummings School of Veterinary Medicine, Tufts University, North Grafton, MA 01536; University of Massachusetts Medical School, Worcester, MA, 01655

**Keywords:** fistulizing Crohn’s disease, microbiome, dysbiosis, perianal fistulas, canine furunculosis

## Abstract

**Background:** Crohn’s disease (CD) is a chronic immune-mediated inflammatory condition caused by the loss of mucosal tolerance towards the commensal microbiota. Approximately 70% of CD patients experience perianal complications. Perianal fistulizing is a predictor of poor long-term outcomes. Animal models of gut inflammation have failed to fully recapitulate the human manifestations of fistulizing CD. Here, we evaluated dogs with spontaneous canine anal furunculosis (CAF), a disease with clinical similarities to fistulizing CD, as a surrogate model for human fistulizing CD.

**Results:** By comparing the gut microbiomes between dogs suffering from CAF (CAF dogs) and healthy dogs, we show that similar to microbiome population trends in CD humans, CAF dogs microbiomes are either very dissimilar (dysbiotic) or similar, yet unique, to healthy dog’s microbiomes. Compared to healthy or healthy-like CAF microbiomes, dysbiotic CAF microbiomes showed an increased abundance of *Bacteroides vulgatus* and *Escherichia coli* and a decreased abundance of *Megamonas* species and *Prevotella copri*. These same determinant bacteria are associated with human CD.

**Conclusions:** Our results highlight the similarities in microbiome community patterns between CAF dogs and CD humans, including overlapping determinant bacterial taxa, and support the use of CAF dogs as a surrogate model to study human fistulizing CD.

## BACKGROUND

Inflammatory bowel diseases (IBD), including Crohn’s disease (CD), are immune-mediated inflammatory condition affecting more than a million individuals in the US. Approximately 70% of CD patients develop strictures or penetrating lesions during their lifetime. Perianal lesions, represent one of the most clinical significant complications of CD, and the presence of perianal fistulizing disease is a predictor of poor long-term outcome [1]. The pathophysiology of perianal fistulizing CD is poorly understood; however, evidence indicates that the gut microbiota is involved [2–4], as treatment with antibiotics (reviewed in [5]), fecal diversion [2, 3], and fecal transplants [4] are beneficial for managing the disease. To our knowledge, no studies have specifically examined the microbiota of patients with perianal fistulizing CD, and there is currently no animal model available for the study of perianal fistulizing CD. Thus, there remains a pressing need for additional studies and a model system to elucidate the contribution of the microbiota to the pathophysiology of perianal fistulizing CD.

We have postulated that canine anal furunculosis (CAF) can be used as a model for studying the pathophysiology and treatment of perianal fistulizing CD [6, 7]. CAF and perianal fistulizing CD exhibit common clinical signs such as inflammation, ulceration of the perianal tissues, and similar cytokine profiles. Moreover, both CAF and CD are likely to develop as a consequence of local T-cell-mediated inflammation [8, 9] and can be treated with immunosuppressive drugs, such as cyclosporin and prednisone [10]. Of note, the German Shepherd Dog (GSD) appears to be overrepresented within the CAF population, with more than 80% of dogs suffering from CAF (CAF dogs) being GSD [11, 12]. GSD are also susceptible to inflammatory bowel disease (IBD) [13, 14], systemic aspergillosis [15–18], and deep pyoderma [19, 20], suggesting that GSD might have a broadly dysfunctional immune response to microbial exposure at epithelial surfaces, similar to human CD patients. Dogs have already proven useful for studying various other spontaneously occurring disorders similar to those affecting humans [6, 21–24], and the dog microbiome is more similar to that of humans than it is to the microbiomes of other animals that are generally used as models of microbiome-centered diseases such as IBD (e.g., mice and pigs) [25]. Dogs also spontaneously develop IBD [13] and can experience common disease complications, including perianal fistulas [26], and exhibit IBD-associated microbiome alterations [27, 28] that resemble changes observed in humans with IBD. Thus, dogs represent an ideal model for studying IBD and in particular perianal fistulizing CD.

Given that the intersection of immunity and the microbiota seems to be at the heart of both perianal fistulizing CD and CAF, we sought to explore naturally occurring CAF as a surrogate model for human perianal fistulizing CD. The aim of this study was to characterize the bacterial microbiome structure of dogs suffering from CAF. We posit that characterizing the microbiomes of CAF dogs will help clarify the pathophysiology of CAF and advise on the translational significance of CAF as an animal model for perianal fistulizing CD.

## RESULTS AND DISCUSSION

### CAF is associated with gut microbiota changes

A total of 20 dogs were recruited for this study: eight healthy control dogs (HC dogs), and 12 CAF dogs (**Table 1**). We collected 4-6 fecal samples per dog, totaling 70 CAF samples and 38 HC samples. Shotgun sequencing of the samples revealed the presence of the following bacterial phyla (relative abundance indicated as percentage of total reads): *Firmicutes* (48%), *Bacteroidetes* (31%), *Proteobacteria* (12%), *Actinobacteria* (7%), and *Fusobacteria* (2%). When we analyzed differences in microbial composition between dogs according to their disease status, we found slight lower alpha diversity in CAF dogs compared to HC dogs (**Figure 1A**, Shannon diversity, Kruskal-Wallis test, *p* value = 0.06), similar to previous results with IBD dogs [29]. As expected, there was also generally lower alpha diversity in older dogs compared to younger dogs (Spearman correlation, *p* value = 0.1). No significant differences in alpha diversity were observed by weight.

**Table 1.** Characteristics of the recruited dogs according to disease status and cluster.

|  | HC dogs<br>(n=8, 38<br>samples) | CAF dogs<br>(n=12, 70<br>samples) | CAF dogs,<br>near cluster<br>(n=7, 65<br>samples) | CAF dogs,<br>far cluster<br>(n=5, 43<br>samples) |
| --- | --- | --- | --- | --- |
| **Dogs description** |  |  |  |  |
| Average age (years) ± STDEV | 7.3 ± 1.43 | 4.3 ± 1.28 | 7.9 ± 1.53 | 7 ± 0.89 |
| Average weight (kg) ± STDEV | 36.9 ± 7.93 | 41.1 ± 7.31 | 42.4 ± 9.37 | 39.2 ± 2.77 |
| Female sex | 4 (50%) | 3 (25%) | 1 (14.3%) | 2 (40%) |
| **Number of fistulas** |  |  |  |  |
| 0-2 | 0 | 4 | 3 | 1 |
| 3-5 | 0 | 5 | 3 | 2 |
| >5 | 0 | 3 | 0 | 3 |
| **Pathology findings** |  |  |  |  |
| *Morphological features* | ND |  |  |  |
| Crypt hyperplasia |  | 3 (25%) | 1 (14.3%) | 2 (40%) |
| Crypt dilation/distortion (% positive) |  | 8 (66.7%) | 5 (71.4%) | 3 (60%) |
| Fibrosis/atrophy (% positive) |  | 8 (66.7%) | 5 (71.4%) | 3 (60%) |
| *Inflammation* | ND |  |  |  |
| Lamina Propria lymphocytes and plasma cells (% positive) |  | 3 (25%) | 1 (14.3%) | 2 (40%) |
| Lamina Propria Eosinophils (% positive) |  | 4 (33.3%) | 2 (28.6%) | 2 (40%) |
| *Final diagnosis* | ND |  |  |  |
| Normal mucosal (% positive) |  | 3 (25%) | 2 (28.6%) | 1 (20%) |
| Lymphoplasmatic inflammatory (% positive) |  | 2 (16.7%) | 0 | 2 (40%) |
| Eosinophilic inflammatory (% positive) |  | 4 (33.3%) | 2 (28.6%) | 2 (40%) |
| Mucosal atrophy/fibrosis (non-inflammatory) (% positive) |  | 5 (41.7%) | 3 (42.9%) | 2 (40%) |

**Figure 1.**
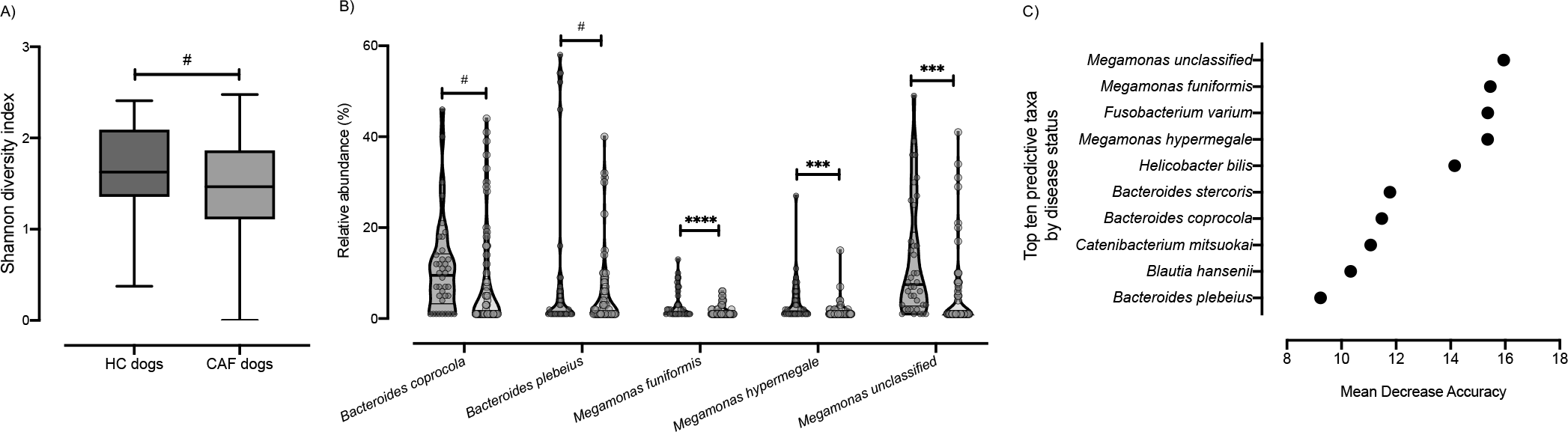
Analysis of the microbiota composition in samples from CAF dogs and HC dogs. A) Shannon index of alpha diversity for HC dogs (green) and CAF dogs (orange). B) Relative abundance of the bacterial taxa found to discriminate between dogs with different disease status: HC dogs (green) and CAF dogs (orange); thick black lines indicate mean abundance. C) Bacterial taxa identified by Random Forest as most strongly distinguishing between samples from dogs with different disease status. FDR t-test, *p* values are expressed as: ****<0.0001, ***<0.001, and #0.1.

After filtering out low abundance genera (i.e., genera not detected four times or more in at least 20% of the samples), we performed gneiss analysis [30], which applies the concept of balance trees to compositional data to identify microbial subcommunities that covary with environmental variables. The gneiss analysis considers the log ratio abundances of subcommunities within the microbiome to indicate taxa whose abundances change relative to other taxa with respect to a variable of interest; in our analysis, the variable of interest was disease status (CAF versus HC). We used linear mixed-effect models to account for inter-individual microbiome variability in this analysis. When we analyzed the microbiomes according to disease status, we found that *Megamonas hypermegale*, *Megamonas unclassified*, *Bacteroides coprocola*, *Megamonas funiformis*, and *Bacteroides plebeius* were differentially abundant in CAF dogs compared to HC dogs (**Table 2**). We further confirmed that there were significant differences in the proportions of these five bacterial taxa between HC dogs and CAF dogs, with lower abundance in CAF dogs (**Figure 1B**, False discovery rate [FDR], t-test). We also applied analysis of composition of microbiomes (ANCOM) to identify discriminant bacterial taxa for CAF dogs versus HC dogs. We identified *M. funiformis* (W = 51), *M. hypermegale* (W = 52), and *M. unclassified* (W = 60) as discriminating between CAF dogs and HC dogs. Altogether, these two robust analyses concurred that the abundance of *Megamonas* species can discriminate between HC dog samples and CAF dog samples.

**Table 2.** Bacterial taxa with significantly different dominance between samples grouped by disease status or cluster.

| Comparison | Determinant bacterial taxa | gneiss, FDR-corrected *p* value |
| --- | --- | --- |
| **Disease status (CAF vs. HC)** | *Bacteroides plebeius* | 0.00137614 |
|  | *Megamonas hypermegale* |  |
|  | *Bacteroides coprocola* |  |
|  | *Megamonas unclassified* |  |
|  | *Megamonas funiformis* |  |
| **CAF dogs in near cluster vs. HC dogs** | *Megamonas funiformis* | 0.01357096 |
|  | *Megamonas unclassified* |  |
| **CAF dogs in near cluster vs. CAF dogs in far cluster** | *Bacteroides plebeius* | 0.00001034 |
|  | *Megamonas hypermegale* |  |
|  | *Bacteroides coprocola* |  |
|  | *Megamonas unclassified* |  |
|  | *Megamonas funiformis* |  |

Lastly, we performed Random Forest analysis to identify bacterial taxa that best distinguished HC dogs from CAF dogs; differences were predicted with an out-of-bag (OOB) error estimate of 7.41%. Akin to what we found above, the top 10 most predictive species belonged to the genera *Megamonas* and *Bacteroides*; however, additional genera, namely *Helicobacter, Fusobacterium, Catenibacterium*, and *Blautia*, were also predictive species of disease status (**Figure 1C**).

### Dogs suffering from CAF exhibit bacterial community structures that resemble those of human CD patients

We next investigated the microbiome patterns further, first using partitioning around medoids with estimation of number of clusters (PAMK) to determine the optimal number of clusters (**Supplementary Figure 1**) and then visualizing the clusters with non-metric multidimensional scaling (NMDS). As seen in **Figure 2A**, CAF dogs were divided into two clusters: healthy-like cluster, referred to hereafter as “near cluster,” gathering in close proximity to HC dogs; and dysbiotic cluster, referred to hereafter as “far cluster,” positioned significantly more distant from the HC dogs (Bray Curtis distance, PERMANOVA, R^2^ = 0.252, *p* value = 0.001). Although the average silhouette width in this analysis indicated a weak structure, this differentiation has been repeatedly reported in patients suffering from CD [27, 31–33]. Taken together, these data suggest that the microbiota of diseased individuals (either CAF dogs or CD humans) can range from one that closely resembles that of a healthy individual to a dysbiotic microbiota that differs markedly from that of a healthy individual.

**Figure 2.**
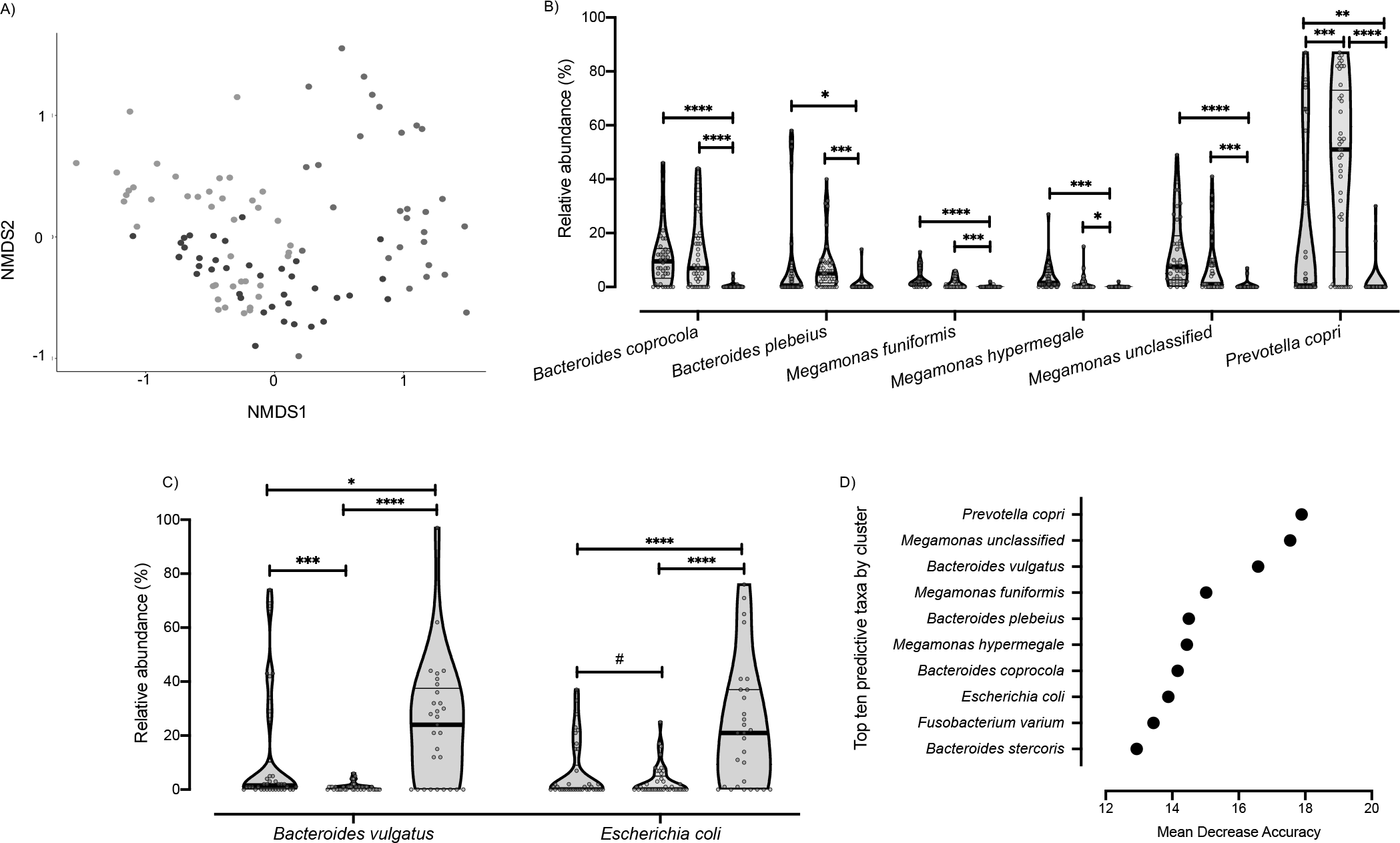
Analysis of the microbiota clusters. A) NMDS visualization of samples from HC and CAF dogs. Bacterial taxa present were quantified by MetaPhlAn, distances were calculated using Bray Curtis dissimilarity index, and samples were plotted based on NMDS. Clusters were defined using PAMK and are colored green (HC dogs), orange (CAF dogs in the near cluster) and red (CAF dogs in the far cluster). B) Relative abundance of the bacterial taxa found to discriminate between dogs, separated by cluster assignment. Clusters are colored as described for panel A, and thick black lines indicate mean abundance. C) Bacterial taxa identified using Random Forest as most strongly distinguishing between samples from dogs belonging to different clusters. FDR t-test, *p* values are expressed as: ****<0.0001, ***<0.001, ** <0.005, * <0.05, and #0.1.

To identify determinant taxa associated with each cluster, we performed additional analyses with gneiss and ANCOM. We found that CAF dogs belonging to the near cluster had different dominance of *M. funiformes* and *Megamonas unclassified* compared to HC dogs (**Table 2**). Moreover, ANCOM results identified *P. copri* (W = 53) as a bacterial taxon that differentiated between CAF dogs belonging to the near cluster and HC dogs.

When comparing CAF dogs belonging to the far cluster and HC dogs, both gneiss and ANCOM identified *B. coprocola* (W = 52), *M. funiformis* (W = 49), *M. hypermegale* (W = 48), and *M. unclassified* (W = 52) as distinguishing between the two groups (**Table 2**). One additional bacterial taxon was identified as being differentially dominant between CAF dogs belonging to the far cluster and HC dogs only by gneiss: *B. plebeius*.

We also investigated differences between CAF dogs belonging to the near and far clusters. We found that the two clusters differed in the dominance of *M. hypermegale*, *M. unclassified*, *B. coprocola*, *M. funiformis*, and *B. plebeius* (**Table 2**). Moreover, ANCOM results identified six bacterial taxa that distinguished between the near and far clusters, specifically, *B. coprocola* (W = 50), *B. plebeius* (W = 51), *Bacteroides vulgatus* (W = 52), *P. copri* (W = 53), *Megamonas unclassified* (W = 48), and *Escherichia coli* (W = 48). Particularly, we found that the relative abundance of six of those bacterial taxa decreased (**Figure 2B**) while two increased (**Figure 2C**) in the far cluster compared to HC dogs and the CAF dogs in the near cluster.

Next, Random Forest was used to identify the most important bacterial taxa within microbial communities for each cluster. With an OOB estimate of error rate of 9.26%, the top 10 predictive bacterial species identified (**Figure 2D**) were similar to those identified using gneiss and ANCOM, as discussed above, and included species belonging to the genera *Megamonas*, *Bacteroides*, *Prevotella*, and *Escherichia*.

Overall, all analyses established that higher abundance of *Megamonas spp.*, *P. copri*, *B. coprocola*, and *B. plebeius* is associated with healthy and healthy-like (near cluster CAF-dog) samples. On the other hand, increased abundance of *E. coli* and *B. vulgatus* is associated with dysbiosis (far cluster CAF dogs).

### Clinical manifestations of CAF are associated with dysbiosis

CAF has a clinical appearance similar to that of perianal fistulas in humans, a complication frequently associated with CD. Specifically, dogs suffering from CAF develop epithelial-lined sinus tracts around the perianal tissue. These ulcerative tracts vary in diameter, depth, and connectivity, and can affect the entire area around the anus. These lesions are usually referred to as fistulas. The CAF dogs recruited in this study presented with varying numbers of fistulas (average, 3.3 ± 1.4. **Table 1**). We found that CAF dogs in the far cluster exhibited a trend of higher average number of fistulas than those in the near cluster (4 vs. 2.87, respectively. **Figure 3A**). Moreover, *Helicobacter canis* dominance was statistically different between CAF dogs according to fistula number, with a negative correlation between *H. canis* abundance and the number of fistulas (**Table 3**).

**Figure 3.**
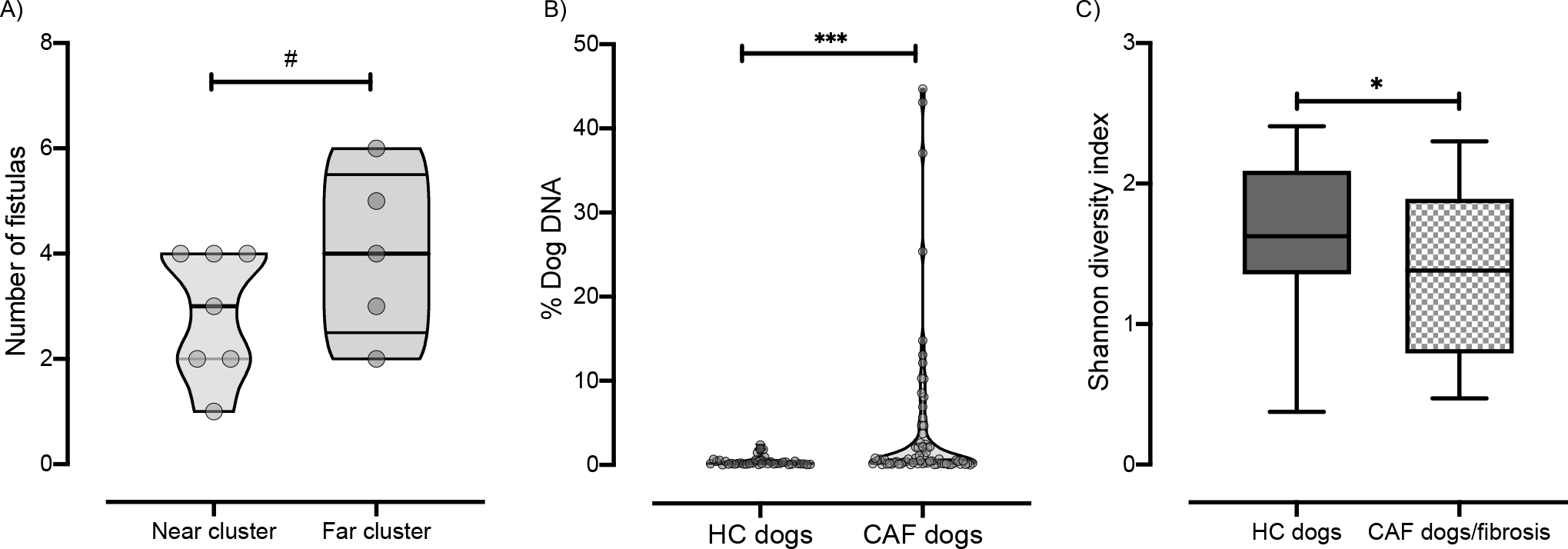
Clinical manifestation of CAF. A) The number of fistulas presented by CAF dogs according to cluster (orange, near cluster; red, far cluster). B) Percentage of dog DNA reads in metagenomic sequencing samples grouped by disease status. C) Shannon index of alpha diversity in samples from HC dogs (green, n = 8) or CAF dogs exhibiting fibrosis (orange pattern, n = 6)

**Table 3.** Bacterial taxa that correlate with clinical manifestations of CAF.

| Disease manifestation/assessment | Determinant bacterial taxa | Spearman correlation | p value | gneiss, FDR-corrected p value |
| --- | --- | --- | --- | --- |
| **Gross examination** |  |  |  |  |
| Number of fistulas | *Helicobacter canis* | -0.4715 | < 0.001 | 0.00000003 |
| **Pathology findings** |  |  |  |  |
| CAF dogs with a combination of crypt hyperplasia, crypt dilation/distortion, and/or fibrosis/atrophy (n = 7) | *Ruminococcus torques* | -0.355 | 0.002 | < 0.05 |
| CAF dogs with a combination of microarchitectural changes and presence of inflammatory markers (n = 5) | *Subdoligranulum unclassified* | -0.1639 | 0.1751 | 0.02305158 |
|  | *Paraprevotella unclassified* | -0.3336 | 0.0048 |  |

It has been suggested that increased host DNA content in fecal samples may be an indicator of inflammation-associated shedding of epithelial or blood cells. Indeed, a previous study found high levels of human DNA in stool samples from pediatric patients with active CD [27]. Thus, we determined the abundance of dog DNA sequences present in each stool sample analyzed as a percentage of total reads. We found that samples from CAF dogs contained a higher percentage (mean = 4.09%, minimum = 0.02%, maximum = 44.68%) of host DNA sequences than did samples from HC dogs (Mean = 0.41%, minimum = 0.02%, maximum = 2.38%, Mann Whitney test, *p* value = 0.001. **Figure 3B**). There was no significant difference between the abundance of host DNA sequences recovered in samples from the near versus far cluster. This result suggests that, compared to their healthy counterparts, CAF dogs have a higher degree of shedding of epithelial or blood cells associated with inflammation, similar to what has been previously observed in CD humans.

In addition to fistulas, GSD with CAF frequently present clinical and histologic evidence of colitis. The major microarchitectural changes accompanying colonic inflammation in dogs include crypt hyperplasia, dilation/distortion, and mucosal atrophy or fibrosis [34]. Thus, we also assessed morphological features and other immune-associated inflammatory markers on the colonic mucosa of the CAF dogs (**Table 1**).

Based on histopathological findings, 75% of the CAF dogs included in this study exhibited morphological evidence of either cryptal hyperplasia, dilation/distortion, or fibrosis. CAF dogs with those major microarchitectural changes showed differential dominance of the same bacterial taxa that were associated with general CAF disease status, namely, *Megamonas* species *(M. hypermegale*, *M. unclassified*, and *M. funiformis)*, *B. plebeius*, and *B. coprocola* (data not shown). In addition to these taxa, *Ruminococcus torques*, known to be decreased in CD [27, 31, 35, 36], was also identified as a determinant taxon between CAF dogs with major microarchitectural changes and HC dogs, with *R. torques* abundance being negatively correlated with microarchitectural changes (**Table 3**). Interestingly, CAF dogs presenting fibrosis (50%) exhibited significantly decreased alpha diversity compared to HC dogs (**Figure 3C**. Shannon diversity, Kruskal-Wallis test, *p* value <0.05). In agreement with our previous analysis according to disease status, CAF dogs exhibiting fibrosis also had lower abundance of *M. unclassified* (ANCOM, W = 48) compared to HC dogs.

Furthermore, CAF dogs exhibiting major microarchitectural combined with infiltration of immune cells (i.e., eosinophils, lymphocytes, or plasma cells) in the lamina propria (41.6%) showed differential abundance of *Subdoligranulum unclassified* and *Paraprevotella unclassified* when compared to CAF dogs without those clinical manifestations. Particularly, those bacterial taxa were negatively correlated with both major microarchitectural changes and immune cell infiltration (**Table 3**). Recently, depletion of *Subdoligranulum* species in dysbiotic CD patients was associated with bile acid dysregulation [33].

Based on the presence of major microarchitectural and immune cells in the lamina propria we calculated pathology scores as described before [34]. We observed that the pathology scores from CAF dogs belonging to the near cluster (median = 2, minimum = 0, maximum = 5) were comparable to those of the far cluster (median = 3, minimum = 0, maximum = 5, Mann Whitney test, *p* value > 0.05).

To investigate whether CAF dogs exhibited peripheral signs of inflammation, complete blood counts (CBCs) were performed. CBC indicators were within the normal range for all tested CAF dogs; thus, no sign of serious inflammation or infection was detectable by CBC analysis (data not shown). Although hemoglobin levels were lower in CAF dogs belonging to the far cluster than in those in the near cluster, we did not identify any bacterial taxa associated with hemoglobin levels, nor with any of the analyzed blood inflammatory markers.

In sum, there is an inverse correlation between *H. canis* abundance and the number of fistulas. In addition to the bacterial taxa associated with disease status, CAF dogs exhibiting microarchitectural changes related to inflammation also exhibit decreased *R. torques* abundance. Furthermore, CAF dogs presenting microarchitectural defects combined with immune cell infiltration in the lamina propria have decreased abundance of *Subdoligranulum unclassified* and *Paraprevotella unclassified*.

### Microbial gene pathways change in accordance with disease status

The genomic content of the microbiota was determined using HUMAnN2 [37]. After filtering for gene pathways that were not present in at least 5% of the samples, we performed clustering analysis on 1,532 filtered pathways. Clustering of the gene pathway abundances separated CAF dogs belonging to either the near cluster or the far cluster from HC dogs (**Figure 4 A**. PERMANOVA, R^2^= 0.19, *p* value = 0.001).

**Figure 4.**
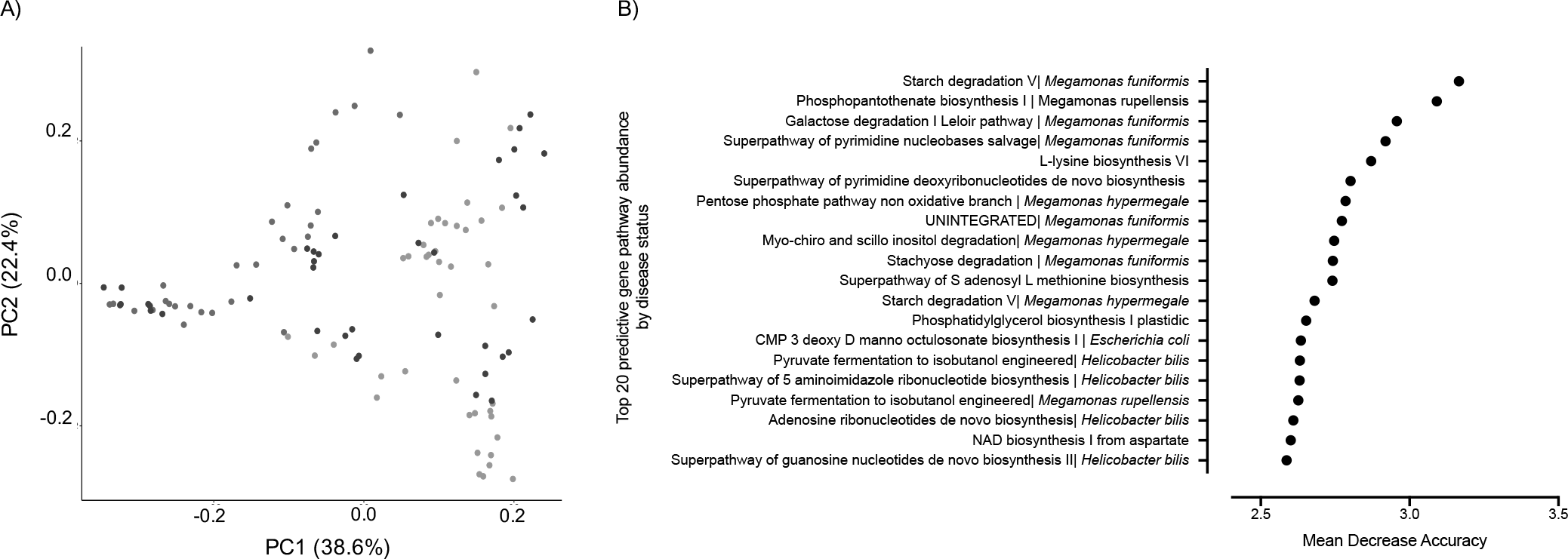
Microbial gene pathway analysis. A) Bray Curtis dissimilarity index beta diversity plot of dogs samples colored by clustering assignation: HC dogs (green) and CAF dogs belonging to the near (orange) or far (red) cluster. Bacterial pathways were quantified by HUMAnN2 and samples were plotted based on principal component analysis. B) The top 20 gene pathways that most strongly distinguish between CAF-dog samples and HC dog samples, identified using Random Forest.

ANCOM results revealed that CAF dogs differed significantly from HC dogs on four of the 1,532 pathways analyzed. The four pathways identified were assigned to the genera *Megamonas* and *Helicobacter* (**Table 4**). Unfortunately, none of the four pathways was attributable to any known pathway in the corresponding database (referred to as “unintegrated”). Comparison of HC dogs with CAF dogs belonging to the near or far cluster revealed 2 or 65 differentially abundant pathways, respectively (**Table 4**). Twelve of these 67 pathways were unintegrated and were assigned to the genera *Helicobacter*, *Peptostreptococcaceae*, *or Collinsella*. Most of the remaining 55 pathways were related to the biosynthesis and degradation of nucleotides, amino acids (particularly branched amino acids), and carbohydrates, the biosynthesis of amines and polyamines, or vitamin K production; these pathways were assigned to the genera *Megamonas*, *Bacteroides*, and *Prevotella*.

**Table 4.** Gene pathways found to be significantly different between samples grouped by disease status or cluster using

| Comparison | Pathway | Bacterial Taxa | ANCOM W value |
| --- | --- | --- | --- |
| CAF dogs vs. HC dogs | UNINTEGRATED | *Megamonas funiformis* | 1528 |
|  | UNINTEGRATED | *Megamonas rupellensis* | 1519 |
|  | UNINTEGRATED | *Megamonas hypermegale* | 1518 |
|  | UNINTEGRATED | *Helicobacter bilis* | 1504 |
| CAF dogs in near cluster vs. HC dogs | UNINTEGRATED | *Megamonas funiformis* | 1453 |
|  | UNINTEGRATED | *Helicobacter bilis* | 1429 |
| CAF dogs in far cluster vs. HC dogs | 1,4-dihydroxy 6-naphthoate biosynthesis II |  | 1338 |
|  | 1,4-dihydroxy 6-naphthoate biosynthesis II | *Megamonas hypermegale* | 1197 |
|  | Adenosine ribonucleotides de novo biosynthesis | *Megamonas hypermegale* | 1325 |
|  | Adenosine ribonucleotides de novo biosynthesis | *Bacteroides coprocola* | 1292 |
|  | Adenine and adenosine salvage III | *Megamonas hypermegale* | 1235 |
|  | Adenosine ribonucleotides de novo biosynthesis | *Megamonas funiformis* | 1225 |
|  | Adenosine ribonucleotides de novo biosynthesis | *Megamonas rupellensis* | 1185 |
|  | Biotin biosynthesis II |  | 1135 |
|  | CDP diacylglycerol biosynthesis I | *Bacteroides coprocola* | 1251 |
|  | CDP diacylglycerol biosynthesis II | *Bacteroides coprocola* | 1251 |
|  | Coenzyme A biosynthesis II mammalian | *Bacteroides coprocola* | 1254 |
|  | D-galactose degradation V Leloir pathway | *Megamonas hypermegale* | 1228 |
|  | D-galactose degradation V Leloir pathway | *Megamonas funiformis* | 1213 |
|  | Folate transformations II | *Bacteroides coprocola* | 1254 |
|  | Folate transformations II |  | 1173 |
|  | Galactose degradation I Leloir pathway | *Megamonas hypermegale* | 1228 |
|  | Galactose degradation I Leloir pathway | *Megamonas funiformis* | 1213 |
|  | GDP mannose biosynthesis |  | 1130 |
|  | Glycolysis IV plant cytosol | *Bacteroides coprocola* | 1320 |
|  | Glycolysis VI metazoan | *Megamonas hypermegale* | 1260 |
|  | Glycolysis VI metazoan | *Megamonas funiformis* | 1186 |
|  | Hexitol fermentation to lactate formate ethanol and acetate | *Megamonas hypermegale* | 1260 |
|  | Hexitol fermentation to lactate formate ethanol and acetate | *Megamonas funiformis* | 1208 |
|  | L-isoleucine biosynthesis I from threonine | *Megamonas hypermegale* | 1271 |
|  | L-isoleucine biosynthesis III | *Megamonas hypermegale* | 1221 |
|  | L-lysine biosynthesis III | *Bacteroides coprocola* | 1261 |
|  | L-lysine biosynthesis VI | *Bacteroides coprocola* | 1257 |
|  | L-ornithine de novo biosynthesis |  | 1160 |
|  | L-valine biosynthesis | *Megamonas hypermegale* | 1300 |
|  | N10 formyl tetrahydrofolate biosynthesis | *Bacteroides coprocola* | 1250 |
|  | Pentose phosphate pathway non-oxidative branch | *Megamonas funiformis* | 1170 |
|  | Pentose phosphate pathway non-oxidative branch | *Megamonas hypermegale* | 1131 |
|  | Peptidoglycan maturation (meso-diaminopimelate containing) | *Megamonas hypermegale* | 1269 |
|  | Phosphopantothenate biosynthesis I | *Bacteroides coprocola* | 1251 |
|  | Phosphopantothenate biosynthesis I | *Megamonas funiformis* | 1140 |
|  | Purine ribonucleosides degradation | *Megamonas hypermegale* | 1256 |
|  | Putrescine biosynthesis IV | *Megamonas hypermegale* | 1252 |
|  | Pyrimidine deoxyribonucleotides biosynthesis |  | 1219 |
|  | Pyrimidine deoxyribonucleotides de novo biosynthesis IV |  | 1212 |
|  | Pyruvate fermentation to isobutanol engineered | *Megamonas hypermegale* | 1207 |
|  | Queuosine biosynthesis |  | 1317 |
|  | Queuosine biosynthesis | *Bacteroides coprocola* | 1224 |
|  | Stachyose degradation | *Megamonas hypermegale* | 1231 |
|  | Stachyose degradation | *Megamonas funiformis* | 1155 |
|  | Starch degradation V | *Megamonas funiformis* | 1276 |
|  | Starch degradation V | *Megamonas hypermegale* | 1249 |
|  | Superpathway of branched amino acid biosynthesis | *Megamonas hypermegale* | 1240 |
|  | Superpathway of glycerol degradation to 13-propanediol |  | 1171 |
|  | Superpathway of L-isoleucine biosynthesis I | *Megamonas hypermegale* | 1193 |
|  | Superpathway of L-threonine biosynthesis | *Megamonas hypermegale* | 1197 |
|  | Superpathway of polyamine biosynthesis I | *Megamonas rupellensis* | 1136 |
|  | Superpathway of polyamine biosynthesis I | *Megamonas funiformis* | 1127 |
|  | Superpathway of pyrimidine deoxyribonucleotides de novo biosynthesis |  | 1405 |
|  | Superpathway of pyrimidine nucleobases salvage | *Megamonas funiformis* | 1324 |
|  | Superpathway of pyrimidine ribonucleosides salvage |  | 1401 |
|  | UMP biosynthesis | *Bacteroides coprocola* | 1244 |
|  | UNINTEGRATED | *Bacteroides coprocola* | 1498 |
|  | UNINTEGRATED | *Megamonas funiformis* | 1495 |
|  | UNINTEGRATED | *Megamonas hypermegale* | 1495 |
|  | UNINTEGRATED | *Megamonas rupellensis* | 1494 |
|  | UNINTEGRATED | *Clostridium hiranonis* | 1390 |
|  | UNINTEGRATED | *Collinsella intestinalis* | 1351 |
|  | Urate biosynthesis inosine 5 phosphate degradation | *Bacteroides coprocola* | 1238 |
| CAF dogs in near cluster vs. CAF dogs in far cluster | Adenosine ribonucleotides de novo biosynthesis | *Prevotella copri* | 1384 |
|  | Guanosine ribonucleotides de novo biosynthesis | *Prevotella copri* | 1376 |
|  | L-lysine biosynthesis III | *Prevotella copri* | 1357 |
|  | L-lysine biosynthesis VI | *Prevotella copri* | 1357 |
|  | Queuosine biosynthesis |  | 1425 |
|  | Queuosine biosynthesis | *Prevotella copri* | 1345 |
|  | UNINTEGRATED | *Prevotella copri* | 1494 |
|  | UNINTEGRATED | *Bacteroides plebeius* | 1493 |
|  | UNINTEGRATED | *Bacteroides coprocola* | 1482 |
|  | UNINTEGRATED | *Megamonas hypermegale* | 1428 |

Random Forest was used to identify microbial gene pathways that best distinguished HC dogs from CAF dogs, predicting differences with an OOB estimate of error rate of 6.48% (**Figure 4B**). As expected, most of the identified gene pathways originated from bacterial genera that were found to be predictive of disease status, specifically, *Megamonas*, *Bacteroides*, *and Helicobacter*.

Altogether, the metabolic capacity of the microbiome varied along with disease status or cluster assignation. Hence, most of the discriminant gene pathways belong exclusively to bacterial taxa depleted in CAF dogs. Although, it is uncertain that this phenomenon causes or is a consequence of disease, characterization of these changes will enlighten our understanding of the microbial dynamics in disease.

## CONCLUSION

Here, we characterized the microbiome of dogs suffering from CAF in order to explore the usefulness of dogs as models for fistulizing CD, a serious complication of CD that represents an unmet clinical need. Our understanding of the pathophysiology of fistulizing CD is not complete albeit their high prevalence and burden for patients with CD. The major challenge to study the pathogenesis of this manifestation is the absence of relevant animal models. Here, we found that dogs suffering from CAF exhibit gut bacterial community structures that resemble those observed in CD patients. Moreover, we found overlapping bacteria species and their metabolic capacity associated with dysbiosis in both CAF and human CD.

Therefore, we propose to use dogs suffering from CAF as a surrogate model to: 1) study fistulizing CD to further elucidate the role of the microbiome in this clinical complication, and 2) explore the pre-clinical efficacy of microbiome-centered therapeutics to treat fistulizing CD.

## METHODS

### Subjects and sample collection

All dogs in this study were GSD that had not received antibiotic treatment within the preceding 3 months. CAF dogs were recruited from Cummings School of Veterinary Medicine at Tufts University. Inclusion criteria included German Shepard dogs with a clinical diagnosis of perianal fistulas with clinical signs of tenesmus, dyschezia, and partial or complete relapse from cyclosporine A therapy. HC dogs were selected based on the absence of perianal fistulas, nonexistence of diarrhea or adverse gastrointestinal symptoms, and lack of immunosuppressant treatment. Dog information including age, weight, gender, number of fistulas, treatment, and antibiotic usage was obtained from clinical records.

Naturally passed feces were collected daily for a period of one week from eight HC dogs and from 12 CAF dogs. Dog owners were provided with tubes (OMNI•gene-GUT| OMR-200, DNAgenotek, Canada) for sample collection and were instructed to place a clean pad on the surface before dogs’ bowel movement to avoid sample contamination from soil, grass, or other surfaces. Samples were freshly collected into the tubes provided. Samples were stored at room temperature until they were delivered at the next clinical appointment, which occurred within 2 weeks of sampling. During that next appointment, biopsies from the distal colon and blood samples were obtained from all CAF dogs for histopathological analysis and CBC, respectively. To determine colonic inflammation, an experienced pathologist blind-scored the biopsies based on the infiltration of eosinophils, lymphocytes, or plasma cells in the lamina propria and morphological changes, including crypt dilation/distortion, fibrosis/atrophy, or crypt hyperplasia, as previously described [38].

### DNA isolation and sequencing

DNA isolation was performed using the MagAttract PowerSoil DNA Kit (Qiagen, # 27100-4-EP) on Eppendorf epMotion 5075 liquid handlers following the manufacturer’s instructions. DNA sequencing libraries were prepared using the Nextera XT DNA Library Preparation Kit (Illumina, #FC-131-1096) and were sequenced on the Illumina NextSeq 500 platform as 150-nt paired-end reads. Read data was quality trimmed and filtered of host DNA using KneadData (https://bitbucket.org/biobakery/kneaddata/wiki/Home) against a prebuilt bowtie2 index for *Canis familiaris* reference genome, build 3.1.

### Metagenomic profiling

Community composition was profiled using MetaPhlan2 [39], and HUMAnN2 [37] was used to assess the content of metabolic and functional genes and metabolic pathways.

### Statistical analysis

We considered only taxa with an abundance ≥0.2% in at least one sample and that were detected in at least 10% of the samples. The Benjamini-Hochberg procedure identified the false discovery rate as 0.25. We used Phyloseq 1.26.1 and the R package cluster v1.4-1 to estimate microbiome patterns using PAMK with optimum average silhouette width [40, 41]. Shannon diversity index (alpha diversity) and Bray Curtis dissimilarity (beta diversity) were calculated using QIIME2 1.8.9. To assess the significance of differences in diversity metrics between samples according to either disease status or cluster assignation, we used Kruskal-Wallis, Spearman correlations, PERMANOVA, ADONIS, and Mantel tests. Determinant bacterial species were identified by the robust algorithms ANCOM [42] and gneiss [30], implemented in QIIME2. For the ANCOM analysis, we analyzed 61 bacterial taxa, and only bacterial taxa reporting W > 45 were considered significant. Separately, we also used ANCOM to identify determinant microbial gene pathways that differentiated between disease or cluster groups; for this analysis, we analyzed 1,532 pathways, and only microbial gene pathways reporting W > 1,100 were considered significant. We also used Random Forest (R package, randomForest 4.6-14), a supervised learning algorithm, to identify bacterial genera that discriminated samples by disease status or cluster assignation. This method identifies specific bacterial taxa or gene pathways that could best classify patient according to a study variable, such as disease status.

## Ethics approval and consent to participate

The protocol for sample collection was approved by the Clinical Research Review Committee of the Cummings School of Veterinary Medicine at Tufts University (CRRC#007-015). Each pet owner consented to participate on the study.

## Availability of supporting data

Sequence files for all samples used in this study have been deposited in the NCBI SRA (SRA: SRP191145 and Bioproject PRJNA53120). Metadata and MetaPhlan tables- with corresponding taxonomic classifications - have been included as Additional files 1 and 2.

## Author contributions

A.M-C, A.H. and L.F. collectively conceptualized the manuscript. A.M-C, D.V.W, J.T, and S.C contributed to dog recruitment, sample management, sequencing and the data analysis. S.C performed colonoscopies and histopathology. A.M-C wrote the manuscript. All authors reviewed and edited the final version of the text.

## Acknowledgments

We thank members of the clinical staff, namely Diane Welsh for her help with dog identification and recruitment for the study.

## Abbreviations

ANCOM: Analysis of Composition of Microbiomes
CD: Crohn’s disease
CAF: canine anal furunculosis
HC: Healthy controls
IBD: Inflammatory Bowel Diseases

---

**Supplementary Figure 1.**
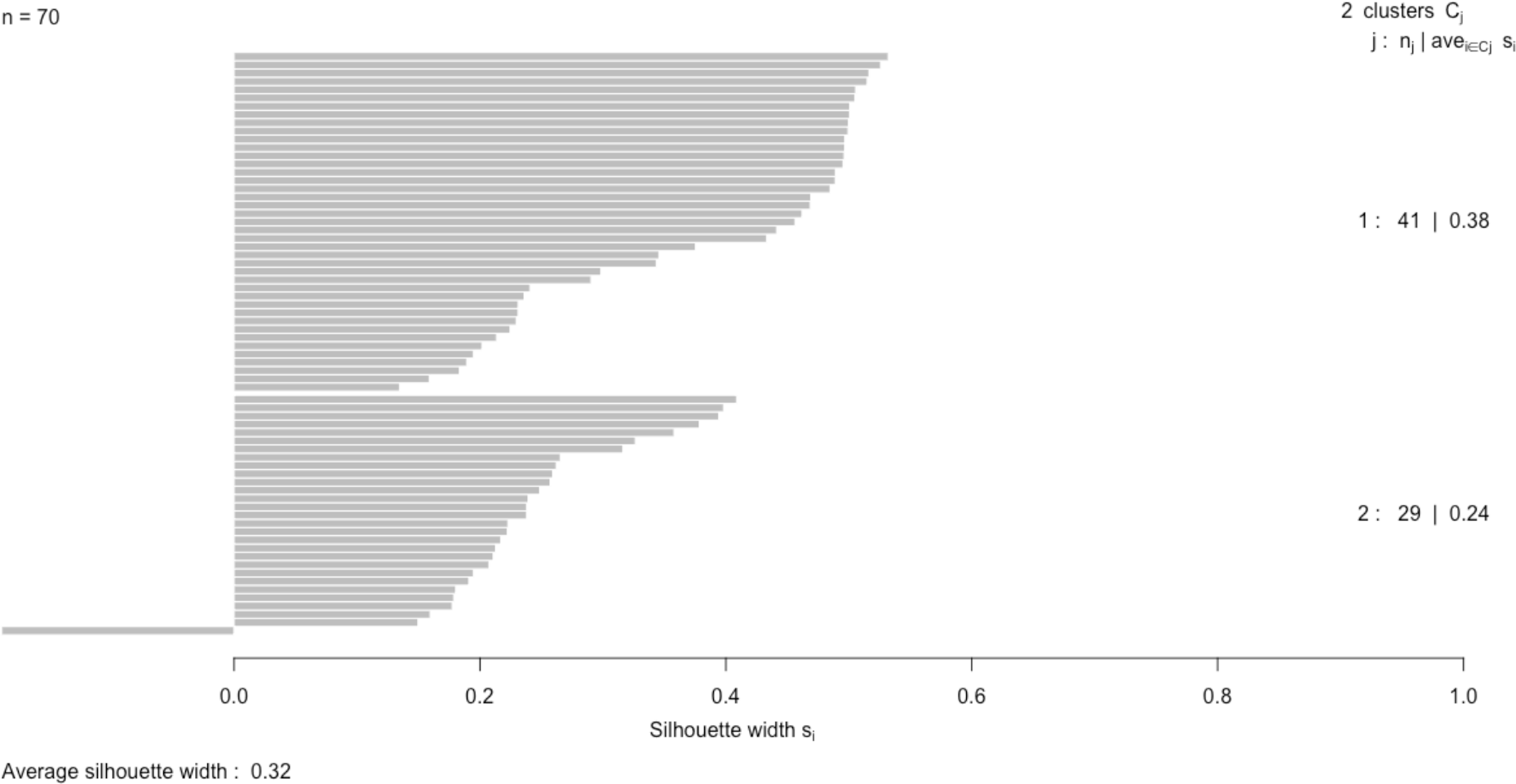
Partitioning around medoids with estimation of number of clusters. Silhouette plot of the clusters identified using the Bray Curtis dissimilarity index.

